# *In Situ* Processing and Efficient Environmental Detection iSPEED of pests and pathogens of trees using point-of-use real-time pcr

**DOI:** 10.1101/871459

**Authors:** Arnaud Capron, Don Stewart, Kelly Hrywkiw, Kiah Allen, Nicolas Feau, Guillaume Bilodeau, Philippe Tanguay, Michel Cusson, Richard C. Hamelin

## Abstract

The increase in global trade is responsible for a surge in foreign invasive species introductions across the world. Early detection and surveillance activities are essential to prevent future invasions. Molecular diagnostics by DNA testing has become an integral part of this process. However, for environmental applications, there is a need for cost-effective and efficient point-of-use DNA testing that would allow for the collection of results in real-time away from laboratory facilities. To achieve this requires the development of simple and fast sample processing and DNA extraction, room-temperature stable reagents and a portable instrument. We conducted a series of tests using a crude buffer-based DNA extraction protocol and lyophilized, pre-made, reactions to address the first two requirements. We chose to demonstrate the use of this approach with organisms that cover a broad spectrum of known undesirable insects and pathogens: the ascomycete *Sphaerulina musiva*, the oomycete *Phytophthora ramorum*, the basidiomycetes *Cronartium ribicola* and *Cronartium comandrae* and the insect *Lymantria dispar*. Tests performed from either infected leaf material or spores (pathogens), or legs and antenna (insects). We were able to obtain positive amplification for the targeted species in all the samples tested. The shelf-life of the lyophilized reactions was assessed, confirming the stability of over a year at room temperature. Finally, successful tests conducted with portable thermocyclers and disposable plastics, demonstrating the suitability of the method, named *in Situ* Processing and Efficient Environment Detection (iSPEED), for field testing. This kit is ideally adapted to field testing as it fits in a backpack and can be carried to remote locations.

## Introduction

The increase in DNA sequences in public databases, driven by advances in sequencing technologies, has revolutionized the molecular diagnostics of pests and pathogens of crops and trees. Molecular identification and detection have become essential components of the prevention, mitigation and management toolbox of forest pests and pathogens [1]. DNA testing allows the rapid, sensitive and accurate detection of target organisms from small amounts of environmental samples harvested during surveys or inspections. This can entirely bypass the time-consuming culture or rearing steps that were previously required to perform a valid identification. The standardization of DNA barcoding databases for fungi and insects [2,3] has generated extensive DNA sequence data that can be exploited to design taxon-specific DNA assays [4–6]. New approaches make use of whole genomes to identify diagnostic genome regions that are translated into highly accurate detection assays [7,8].

The Polymerase Chain Reaction (PCR) is a powerful method to amplify DNA fragments. The development of instruments that can measure fluorescence in real-time [9] allowed the development of single-step DNA detection assays, removing the need to visualize the PCR product by gel electrophoresis based on fragment size. The use of fluorescent DNA binding dyes such as SYBR green [10] or the use of internal probes labelled with a dye [11] enables more complex assay creation. For example, multiplexing can be achieved by using different dyes on probes targeting different genes or organisms. There are now thousands of PCR assays for the detection of pests and pathogens [12,13], and several are used operationally [14].

One drawback of PCR-based detection until recently is that it needs to be conducted in a laboratory environment, with large benches and sensitive instruments (thermal cyclers, centrifuges, pipettes) to handle the small volumes of reagents. Most chemicals and reagents used in the reactions are typically not stable at room temperature for long periods and must be maintained at 4°C or frozen. DNA extraction methods generally require multiple steps and non-trivial equipment. They usually comprise a mechanical disruption step, often in a homogenizer, followed by lysis of the sample. DNA is then purified from the lysate using liquid-liquid extraction or bound to a silica gel matrix before washing and elution. These processes are time-consuming and costly and require multiple buffers for lysis, washes and elution as well as several instruments: water bath, centrifuge and several pipettes for different volumes. Similarly, real-time PCR reagents typically require storage at −20°C (Quantifast Probe and Quantitect Probe from Qiagen or SsoAdvanced Universal Probes and iTaq Universal Probes from Bio-Rad for example). Furthermore, the reaction setup involves pipetting microliter-scale volumes multiple times.

Having the capacity to perform PCR on-site would be beneficial for environmental applications that must be performed in the field. However, the complexity of the extraction and amplification processes and the cost of sample processing have hampered the development of such PCR testing, increasing turn-around time to obtain results. This can be a disadvantage for the detection and identification of forest invasive pests and pathogens because sampling is often conducted in remote locations. Sending samples to a laboratory and obtaining results often requires several days. Field-based DNA testing would speed up the process and could inform additional sampling strategies, for example, if the presence of a threatening pest or pathogen is discovered. This could trigger more rapid mitigation and management actions, a crucial aspect of prevention [15].

In recent years, medical point-of-care and military point-of-need applications of DNA tests have generated interest in designing small real-time PCR instruments, ranging from transportable, such as the Coyote Bioscience Mini8 Plus (Zhang et al., 2017) and the Bio Molecular Systems Mic (Valtonen et al., 2019) to handheld such as the Biomeme two3 and Franklin (Russell et al., 2018). Herein we describe the development of a complete sample-to-data, point-of-use, real-time PCR system to provide *in situ* Processing and Efficient Environmental Detection (iSPEED) of pests and pathogens. This system can easily be carried in a backpack and is ideal for environmental applications. It comprises a minimal DNA extraction method to process environmental samples without centrifuge or homogenizer instruments and room-temperature stable ready-to-use real-time PCR reagents that we used in portable instruments. We demonstrate a broad range of portable applications targeting four forest pests and pathogens that are a concern for forest health.

## Material & Methods

### Plant, fungal and insect material

All material was obtained as part of surveys of tree diseases or insect pests or via artificial inoculation of plant material. The number of extractions performed or tests conducted are mentioned in the extraction or results sections. We tested direct PCR detection of pathogens on leaves and stems of naturally-infected poplars and pines. Poplar leaves infected with *Sphaerulina musiva*, causal agent of the Septoria poplar leaf spot and canker, were collected from a plantation in Salmon Arm, British Columbia in 2015 as described previously [16]. A leaf disk (about 4 mg) was cut out from a poplar hybrid (*P. trichocarpa* x *P. deltoides*) leaf infected with the pathogen *S. musiva* and further cut in two halves (Fig 1A); each half was used for comparison of two DNA extraction methods (described below).

**Figure 1:**
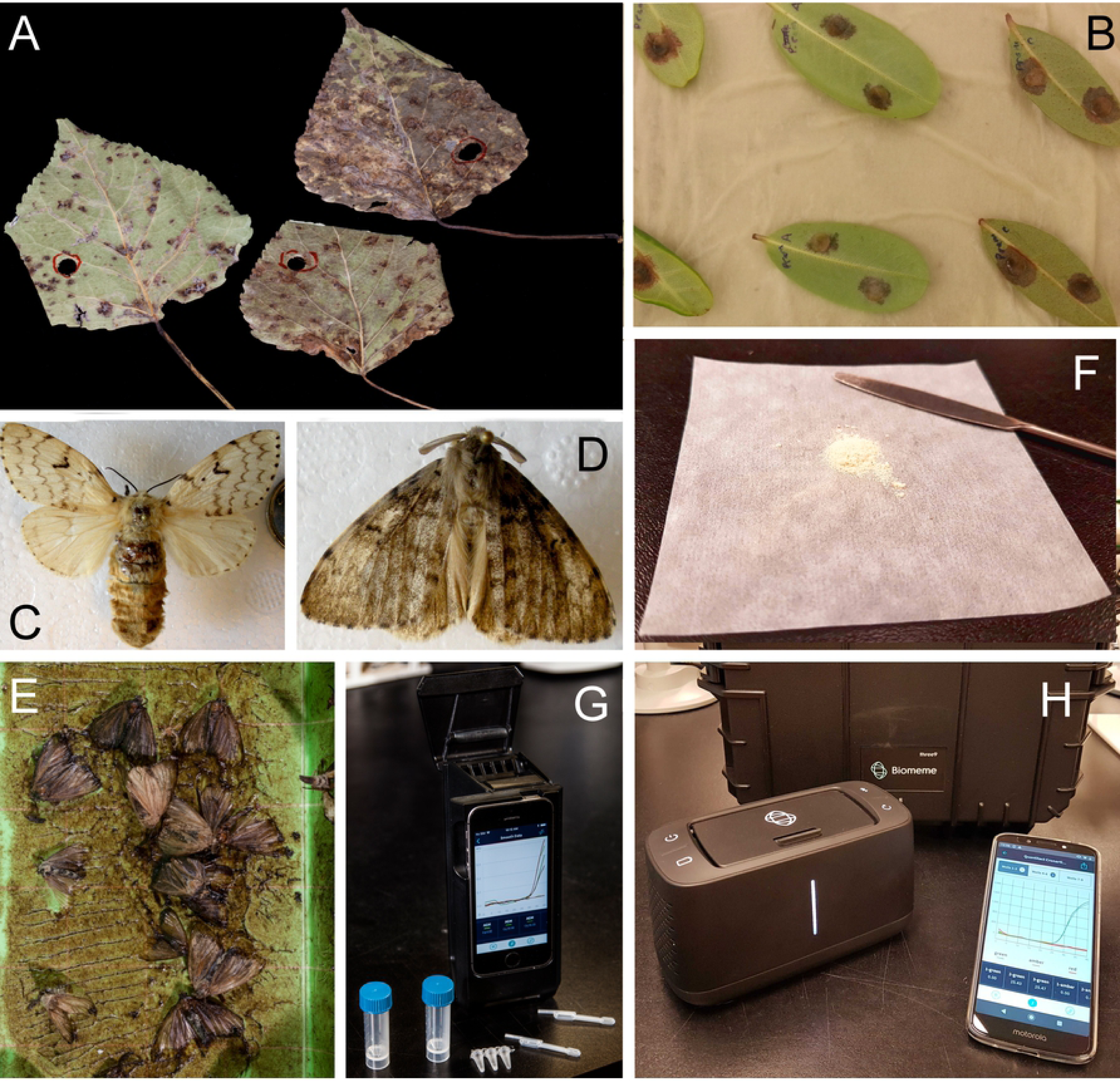
Sample-to-data point-of-use real-time PCR system to provide *in situ* Processing and Efficient Environmental Detection (iSPEED) of pests and pathogens: A) *Sphaerulina musiva* infected poplar leaves; B) *Phytophthora ramorum* infected Rhododendron leaves; C) *Lymantria dispar dispar;* D) *Lymantria dispar asiatica*; E) An example of the moth specimens collected in a pheromone trap; F) *Cronartium ribicola* aeciospores; G) The two3, a first-generation hand-held real-time PCR instrument, with the plastic consumables used for field-trials: 5 mL screw cap vials, a PCR strip and 20 µL pastettes; H) The Franklin, a portable real-time PCR instrument used for the Point-Of-Use field-assay, along with a smartphone used to set-up the experiments, check the results and upload them to a remote server.

Aeciospores of the Comandra blister rust fungus of pines, *Cronartium comandrae*, were collected in May 2018 from stems and branches of lodgepole pines (*Pinus contorta*) in Smithers, British Columbia and stored at 4°C at 31% relative humidity for 7 days before being transferred to −20°C. Aeciospores of the white pine blister rust fungus, *Cronartium ribicola*, were collected in April 2019 from Cypress Mountain, British Columbia and kept at 4°C until used. Approximately 2.5 mg of spores were used per extraction.

For *Phytophthora ramorum*, the causal agent of sudden oak and sudden larch death, we inoculated leaves of *Rhododendron macrophyllum, R. purdomii* and *Rhododendron spp.* collected at the University of British Columbia Botanical Garden. A 0.5-1.0 cm surface incision was cut on the abaxial surface of surface-sterilized detached leaves with a scalpel without puncturing the adaxial side of the leaf. A 10-day-old agar plug of a *P. ramorum* culture of the NA2 lineage (04-38813; Gagnon et al., 2017) grown on carrot agar at 23°C in the dark (Leslie and Summerell, 2006) was placed mycelium side face down over the wound. Inoculated leaves were placed on damp kimwipes in a metal tray and enclosed in a plastic bag with a breathable patch to maintain air circulation and humidity. The leaves were incubated in the dark for 6 days in high humidity at 19-21°C, after which a 5 mm leaf disc was cut from the edge of the necrotic zone (about 20 mg). The leaf disc was cut in half then each half used in the different extraction methods. We also used DNA extracted from cultured mycelia NA1 PR-11-010, described in Feau *et al.* [18], as a positive control.

Pinned insects belonging to the genus *Lymantria* were obtained from collectors from Lorgues, France, in 2009 and from Krutinka in the Omsk Oblast, Russia, in 2013 [6]. In addition, samples were obtained from *Lymantria* pheromone traps deployed in Vancouver, BC, in 2016 as part of the annual gypsy moth survey by the Canadian Food Inspection Agency (CFIA). Either a single leg or a pair of antennae were used for extraction (1-2.5 mg of material).

### DNA extraction

DNA was extracted using established protocols as well as using a field-ready protocol. DNA was extracted from *S. musiva* lesions cut out from infected poplar leaves and from *Cronartium* spores using column-based DNeasy Plant Mini (Qiagen, Venlo, Netherlands) following the manufacturer’s instructions, modified as such: disruption was performed by grinding using three 3 mm steel beads in a MM300 mill (Retsch, Haan, Germany) for 2 minutes at 30 Hz. For *P. ramorum* cultures, total genomic DNA was extracted by the CTAB (cetyl trimethylammonium bromide) method. DNA was eluted in TE (Tris –EDTA) buffer (10 mM Tris-HCL, 0.1 mM EDTA, pH 8) and used as template [19]. DNA was extracted from single or multiple antennae or legs of *Lymantria* using the DNeasy Blood & Tissue kit (Qiagen, Venlo, Netherlands) following the manufacturer’s insect protocol. Disruption was performed by grinding using a single 5 mm steel bead in a MM300 mill (Retsch, Haan, Germany) for 2 minutes at 30 Hz.

A simplified field-ready cost-effective DNA extraction method was developed by using Edwards buffer [20]: Tris 200 mM, EDTA 25 mM, NaCl 250 mM, SDS 0.5% (w/v). For the naturally infected poplar and the inoculated Rhododendron leaves, 1% PVPP (w/v) (Millipore Sigma, Darmstadt, Germany) was added to the buffer to mitigate the effect of phenolic compounds prevent in leaves on the PCR reactions. Infected plant or insect material was incubated in 40 µL of buffer for 10 min at 95°C to release the nucleic acid into the buffer solution. The extract was then used as template for PCR either as a 50-fold or 100-fold dilution in water, depending on the tissue.

### Real-Time PCR TaqMan assays

#### Real-time PCR assays

We used real-time PCR assays that have been previously developed, tested and validated for *Sphaerulina musiva, P. ramorum* and European gypsy moth (EGM) Lymantria (Table S1). The *S. musiva* assay used the Genome-Enhanced Detection and Identification (GEDI [8]) approach and amplifies a genome fragment (SepMu) found in *S. musiva* but in none of the close relatives and also a fragment of the RuBisCo large subunit gene from the plant DNA (RbcL), as an internal control [16]. The *P. ramorum* assays were TAIGA-C62, also identified using the GEDI approach, targeting a single copy nuclear region and providing high levels of specificity [18] and TrnM, which amplifies a portion of the mitochondrial genome, giving the assay a higher sensitivity [21]. The gypsy moth assay is a duplex real-time PCR that can distinguish between the asian and european allele of the FS1 marker [6].

For the rust fungi, we designed an assay that can differentiate the native *C. comandra* blister rust from the invasive exotic *C. ribicola* pine rusts and their hybrids [22]. The assay targets the glutamine synthetase gene (*Dcon10*) from *C. ribicola* and *C. comandrae* (NCBI PopSet 63080856). The primers and probes were designed using Primer3 [23,24] and IDT oligoanalyzer (IDT, Skokie, IL, USA). A TaqMan probe using LNA bases (to increase specificity) was designed for each of *C. ribicola* and *C. comandrae* (Table S1).

#### Laboratory real-time PCR testing using fresh reagents

Reactions performed in the lab were conducted with the Qiagen QuantiTect Multiplex PCR mastermix (Qiagen, Venlo, Netherlands) following the manufacturer’s instructions in 20 µL. The instrument used was an Applied Biosystems Viia7 thermocycler (ThermoFisher Scientific, Waltham, MA, USA). Cycling parameters were the following: 15 min at 95°C, 40 cycles of 15 sec at 95°C, 90 sec at 60°C, as specified by the manufacturer. Details of the reaction set-up for each assay can be found in Table S2.

#### Field-ready mastermixes

Field-ready reaction mixes used lyophilized reagents that comprised the QuantiTect mix. Primers and probes were added to 10 µL of the mix from the 100 µM stocks, and trehalose was added to a final concentration of 5% from a 30% stock to increase the stability of the reagents. The reactions were aliquoted in 100 µL MicroAmp Fast 8-tube strips (ThermoFisher Scientific, Whaltam, MA, USA), frozen at −20°C and then lyophilized for 60 to 90 minutes in a Freezone 2.5 Liter freeze-drier (Labconco, Kansas City, MO, USA). After lyophilization, the strips were stored in the dark, either at room temperature or −20°C. To conduct the real-time PCR assays, the lyophilized reagents were resuspended in a final volume of 20 µL. For reactions performed with DNA extracted with kits, two microliters of DNA were diluted in 18 µL of water. The reactions prepared using the direct PCR method were resuspended in 20 µL of a 1/50 or 1/100 dilution of the crude DNA extract. The stability of the lyophilized reagents was tested every three months by running real-time PCR with the tubes that were stored frozen and those stored at ambient temperature. Detail about each assay is provided in Table S3.

#### Field-ready extraction and real-time PCR using portable instruments

Field-ready real-time PCR testing was conducted on portable real-Time PCR thermocyclers (Biomeme, Philadelphia, PA, USA). The DNA extraction and real-time PCR reactions were performed with 20 μl disposable pipettes (pastettes; Alpha Laboratories, Eastleigh, UK) that were used for the dilution of the extractions and the pipetting of the samples into the real-time PCR strips.

## Results

### Assembling a real-time PCR field kit: DNA extraction and real-time PCR reagents

Crude DNA extraction methods that can be used in the field should be simple and require no instrument. For environmental applications, DNA extraction should be cost-effective. Methods have been developed previously for blood (Mercier et al., 1990), fixed tissue samples (Panaccio et al., 1993), insects (Czank, 1996) and fungi (de los Ríos et al., 2000) among others. Our field-based crude DNA extraction method involves a simple protocol that does not require instruments and uses a basic inexpensive Tris buffer (Edwards buffer) containing SDS (Edwards et al., 1991). It was chosen due to its effectiveness, simplicity and very low-cost.

### Efficacy of field-ready DNA extraction method on *Sphaerulina musiva* poplar pathogen

As a first proof of concept experiment, we compared the efficacy of our field-ready DNA extraction method against a column-based DNA extraction. We chose an assay that can detect *Sphaerulina musiva*, a fungal pathogen that causes leaf spot disease (Fig 1A) and perennial stem and branch cankers in poplars [25]. While relatively harmless in natural forests, *S. musiva* is a significant threat to hybrid poplars grown in plantations in North America [26,27]. The leaf spot symptom caused by *S. musiva* is impossible to differentiate from those caused by the related species *S. populicola*, a foliar pathogen that does not cause cankers [28]. This pathogen has recently established itself in new regions, including western North America, where it threatens poplar plantations (Herath et al. 2016).

Both the field-ready and the column-based extractions yielded a positive real-time PCR reaction from infected leaf spots for the internal leaf control and the pathogen assay. Although the cycle threshold (C_t_) values were lower for the column-based extraction DNA (ANOVA: F = 119.58, p< 0.001) the field-ready extraction using the Edwards buffer extraction yielded unambiguously positive detection of the pathogen, with C_t_ values ranging from 21.24 to 25.41 (Fig 2A, Table S4). Those C_t_ values are comparable to those observed in another study using extraction kits [16]. The higher C_t_ values in the field-ready extractions compared to the column-extracted DNA were expected given the thorough disruption used before the column-based extraction and the likely presence of inhibitors in the crude extract.

**Figure 2.**
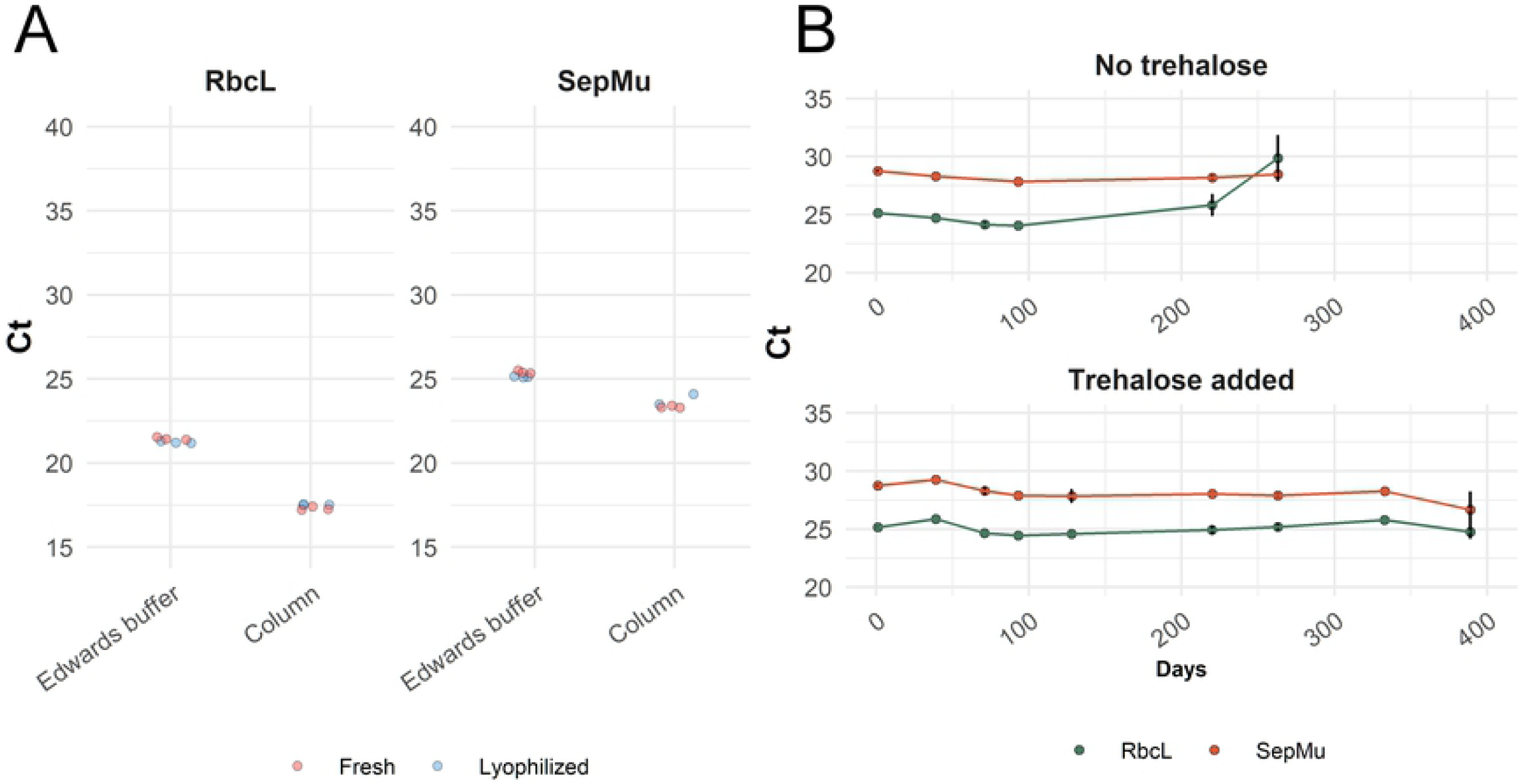
Real-time PCR detection of *Sphaerulina musiva* from naturally-infected poplar leaves using field-ready and laboratory protocols. A) C_t_ values obtained using the plant control RbcL (left) and *S. musiva* assays (right), comparing the fresh vs lyophilized PCR reagents and the two extraction methods; B) shelf-life test using lyophilized reactions with or without trehalose added.

To be able to use this real-time PCR assay in the field, reagents needed to be stable at ambient temperature. We pre-combined all required PCR reagents so that only the sample extracts diluted in water need to be added. Initial trials using lyophilized real-time PCR mastermixes containing the polymerase, reaction buffer, dNTP, primers and probe showed that the reagents could tolerate the process (data not shown). Strip PCR tubes were prepared with the same real-time PCR mastermix with trehalose added to reach 5% of the reaction volume. Trehalose is a cryoprotectant [29] as well as a PCR enhancer [30]. This field-ready approach was tested with the *S. musiva* assay - detection of the plant material and the pathogen was achieved in all reactions, with C_t_ values of 17.5 and 23.8 using column extracted with the plant DNA, for RbcL and SepMu, respectively (Fig 2A, Table S4). The lyophilization was not a significant factor when comparing the C_t_ values of freshly prepared and freeze-dried reactions (ANOVA: F = 0.05, p>0.9), indicating that the lyophilized reactions were performing adequately.

### Detection of *Phytophthora ramorum* on infected rhododendron leaves

We tested the applicability of this approach to the oomycete *Phytophthora ramorum*, an exotic plant pathogen under quarantine or regulated in multiple countries, including Canada [31], the USA [32] and the European Union [33]. *P. ramorum* has been causing large, deadly, epidemics on oaks in the USA [34] and larch in the UK [35] and is capable of infecting a wide range of hosts, including many ornamental plants [36]. Rhododendron is a common host for *P. ramorum* and a frequent carrier of the pathogen in nurseries, making it an ideal candidate to test the effectiveness of the method for *P. ramorum* detection *in planta*.

We compared DNA detection by real-time PCR for the *P. ramorum*-infected leaf discs using either the DNA extracted with the field-ready Edwards buffer (1:100 dilution) with direct real-time PCR or for pure cultures with the columns. For the field-ready extraction, we placed half of the leaf disk in the Edwards buffer and heated it at 95°C for 5 min before conducting real-time PCR with fresh reagents; we used the other half of the leaf disk to perform real-time PCR with the lyophilized reagents. DNA from the pure culture and all hosts inoculated with *P. ramorum* produced a detectable amplification signal, with C_t_ values ranging from 15.77 to 32.16 for the TrnM mitochondrial assay and from 21.93 to 34.62 for the TAIGA-C62 nuclear assay; none of the uninoculated leaves yielded detectable signals (Fig. 3, Table S5). All combinations of field-ready Edwards buffer extractions and lyophilized reactions yielded detectable signal, although fresh reactions yielded slightly lower C_t_ values than lyophilized reactions (F = 28.43, p<0.0001). Cycle threshold values for the TrnM mitochondrial assay were lower than for the TAIGA-C62 nuclear assay (F = 316.18, p<0.0001). This result is expected because of the higher sensitivity of the multi-copy mitochondrial assay than the single-copy nuclear assay (Feau et al., 2019; Bilodeau et al., 2014). Since there were no false positives and no false negatives in our testing, this experiment provides support for the suitability of this approach for fast and accurate detection and identification of this pathogen and could become a core tool in the management, mitigation, and containment of current infection centers and in preventing new invasions [37].

**Figure 3.**
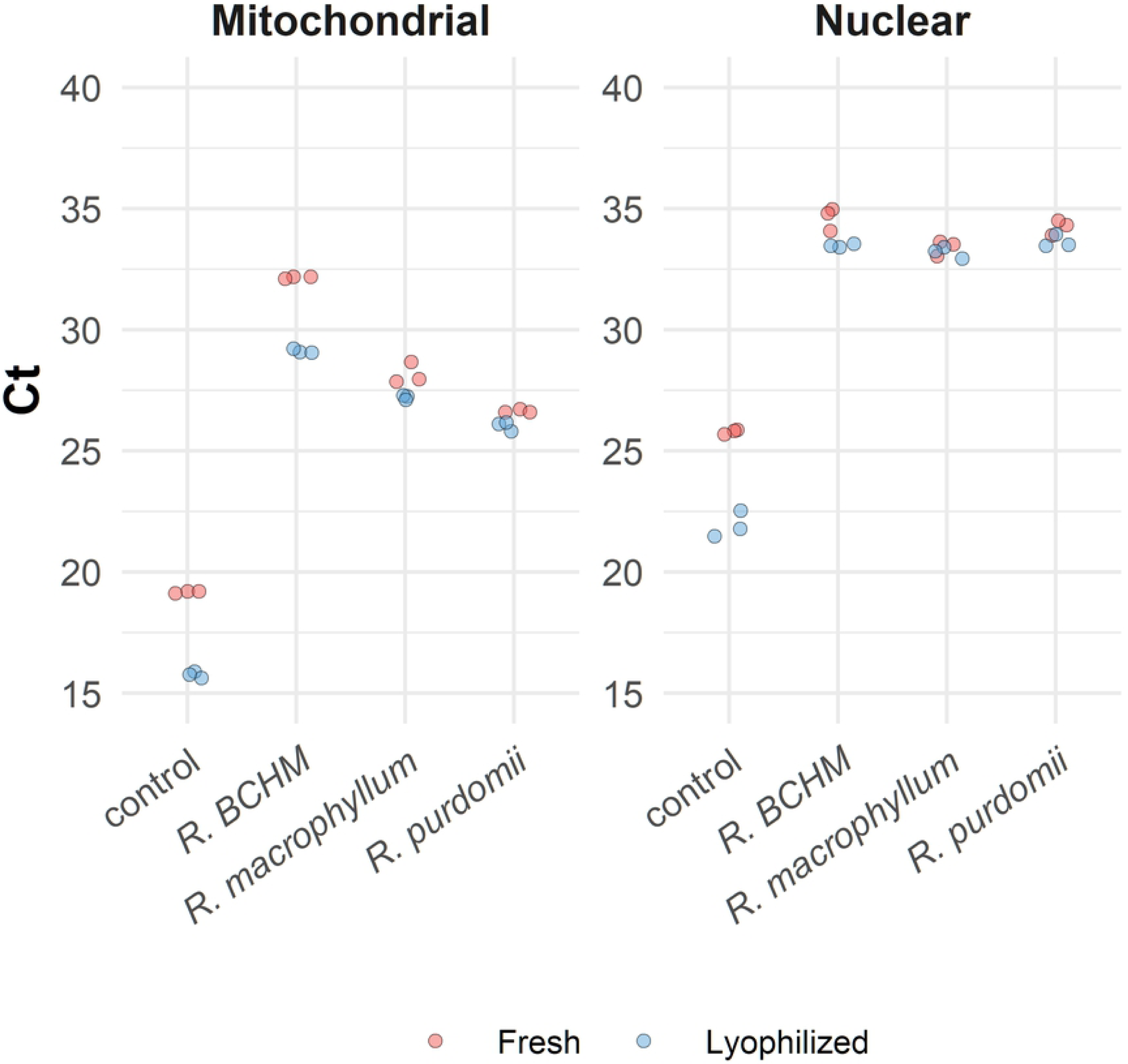
Real-time PCR amplification of *Phytophthora ramorum* from artificially infected *Rhododendron* leaves using field-ready and laboratory protocols. Cycle threshold values obtained with Rhododendron leaves from different species infected with *P. ramorum* as well as DNA extracted from a cultured sample (control). The figure contrasts fresh/lyophilized results. The Mitochondrial assay (left) targets a *P. ramorum* mitochondrial DNA sequence while the nuclear assay (right) targets a single copy *P. ramorum* nuclear locus

### Identification of Asian and European gypsy moth

We wanted to demonstrate that our approach can be applied to insects as well. Molecular identification of invasive insects through real-time PCR has been used successfully applied to multiple species, such as fire ants (*Solenopsis invicta*), the small hive beetle *Aethina tumida* and several mosquito species (*Aedes spp.*) [38–40]. We tested our protocols on gypsy moths (*Lymantria spp.*), defoliators that produce caterpillars able to feed on multiple tree species, especially oaks (*Quercus spp.*), trembling aspens (*Populus tremuloides*) and birches (*Betula* spp.) [41]. The European gypsy moth (EGM), *Lymantria dispar dispar*, is established in eastern North-America [42]. The Asian gypsy moth (AGM), *Lymantria dispar asiatica*, is present in several countries in Asia and has been found several times in North America but has not established itself. Efforts to prevent its introduction and establishment require conducting surveys and inspections which, in turn, require reliable identification. One of the concerns is that AGM as a broader host range than EGM that includes conifers and their females can fly up to 25 km, whereas EGM females are flightless. AGM and EGM can hybridize and could produce flight-capable hybrids that may escape identification [43]. Morphological identification of adults and larvae of the two species is difficult at best and impossible for eggs. A series of gypsy moth TaqMan assays that can together identify *L. dispar asiatica* from several close species such as *L. dispar dispar, L. postalba* and *L. albescens*, was recently published [6].

The AGM4 duplex real-time PCR assay targets two alleles of the nuclear marker FS1. The major alleles are different in Asian Lymantria populations and North-American populations [6]. We assessed the performance of our field-ready protocol using this multiplex assay and tested it using pinned and trapped moth specimens. DNA was extracted with the field-ready extraction as well as the column-based DNeasy Blood and Tissue kit (Qiagen, Venlo, Netherlands) from a single leg of pinned specimens (Fig1C and D). The EGM individual was positive for the FS1 North American allele using lyophilized and fresh real-time PCR reagents but was negative for the Asian allele (Fig 4, Table S6). In contrast, the AGM sample was positive for the Asian FS1 allele and negative for the European allele using both column and field-ready extractions and fresh as well as lyophilized reagents (Fig 4, Table S6). An ANOVA showed no significant difference between fresh and lyophilized reactions or between the extraction methods (F = 3.770, p>0.05).

**Figure 4:**
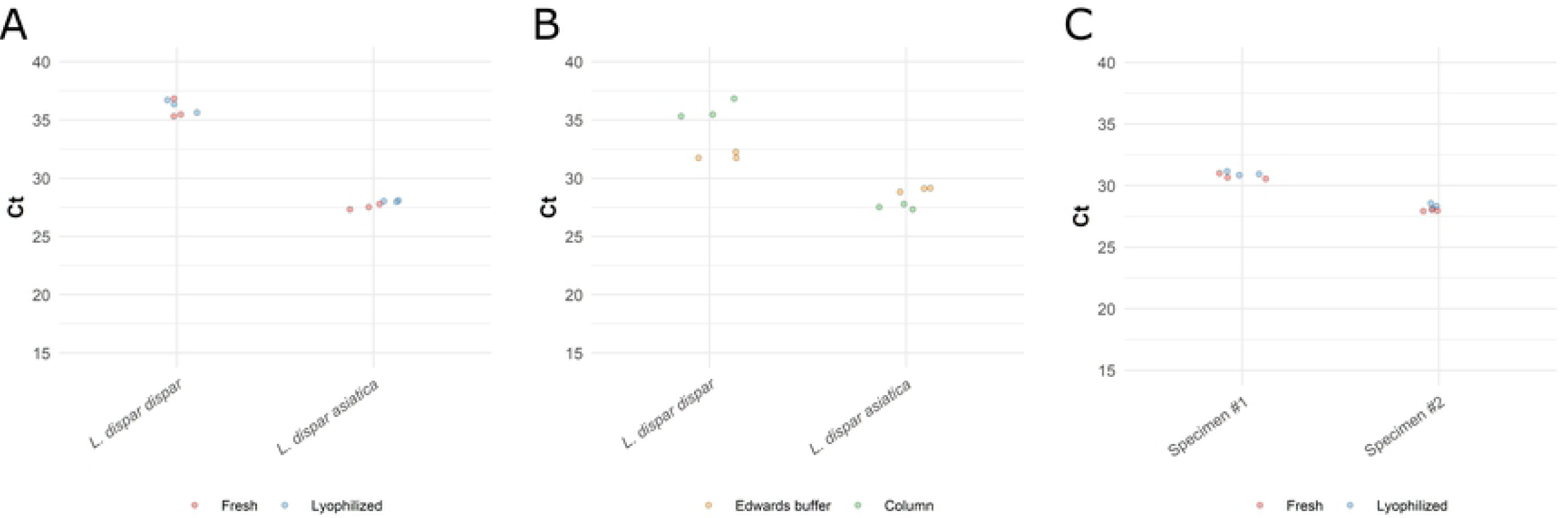
Real-time PCR detection of *Lymantria dispar* using field-ready and laboratory protocols. The C_t_ values shown on panel A and B are for the European FS1 allele for *L. dispar dispar* and the Asian FS1 allele for *L. dispar asiatica*. A) Results of the tests using column extracted DNA with fresh and lyophilized reactions. B) Results of the test comparing column extracted vs. field-ready DNA extraction reporting C_t_ values for the same individuals. C) Results of the test using samples collected from a pheromone trap, with fresh and lyophilized reactions, the C_t_ values shown are for the European FS1 allele.

We tested our protocols with insects obtained from pheromone traps (Fig 1E, Table S6). Typically, those traps remain in the field for several weeks and contain tens to hundreds of insects that, in North America, are presumed to be EGM. We used the field-ready extraction by placing a pair of antennae for each of two specimens in the Edwards buffer and heating them at 95C for 5 min before conducting real-time PCR with both fresh and lyophilized reactions. Both specimens tested amplified only the North-American FS1 allele (Fig 4C, Table S6). An ANOVA showed a small but significant difference between fresh and lyophilized reactions (F = 9.236, p<0.05). These experiments demonstrate that the combination of field-ready DNA extraction and reagents can be applied to insects and provide rapid and accurate identification of the Asian and European subspecies of Lymantria.

### *Cronartium* blister rust pathogen of pine

We wanted to test the use of our field-ready approach directly on fungal spores. Rust fungi of the *Cronartium* genus are basidiomycetes comprising pathogens that cause cankers and blisters on the stems and branches of pines. *Cronartium ribicola* is responsible for the white pine blister rust disease that contributed to the decline of North American populations of five needle pines and the listing of whitebark pine as an endangered species (Fig 1F) [44]. *Cronartium comandrae* can attack two and three needle pines such as *Pinus banksiana, Pinus contorta* and *Pinus ponderosa* and can also cause mortality in these pines that are widely distributed in Canada and of are of commercial importance for lumber and pulp production [45]. The discovery of new alternate hosts for *C. ribicola* and of hybrids between *C. ribicola* and *C. comandrae* has created renewed efforts to monitor these pine pathogens [22,46].

The assays specific to *C. ribicola* and *C. comandrae* detected these pathogens in all the tests with C_t_ values ranging from 26.5 to 32.9, indicating the suitability of the combination of field-ready extraction and lyophilized reactions for use directly on spores of rust fungi (Fig 5A, Table S7). The different samples and reagent conditions did not yield significantly different C_t_ values for the *C. comandra* samples, but the extraction methods were significantly different (F = 61.1248, p<0.001); similarly, fresh and lyophilized reactions did not impact Ct values (F = 0.333, p>0.3) in *C. ribicola*, but extraction methods and extraction batch significantly affected C_t_ values in *C. ribicola* (F = 273.166, p<0.0001 for method and F = 125.214, p<0.0001 for extraction number); this is possibly a reflection of the variation in spore abundance in each blister. Although the thorough mechanical disruption and purification provided by the column-based DNA extraction improves the performance of the DNA for real-time PCR, we found no false negatives, indicating the potential for field application of the method.

**Figure 5.**
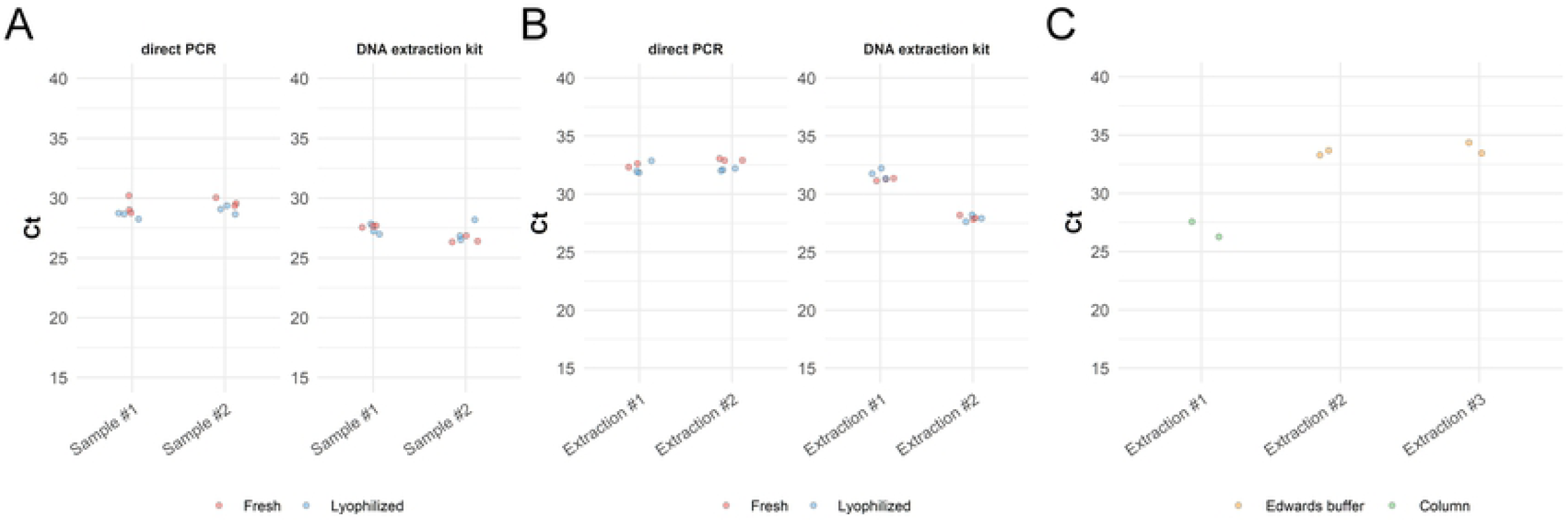
Real-time PCR amplification of *Cronartium spp.* spores collected on pines. Average C_t_ values obtained with *Cronartium* spores, contrasting fresh and lyophilized and field-ready extraction and kit results. A) Results using spores from *C. comandrae*. Spores from two different blisters were used, labelled as sample #1 and sample #2. DNA from spores from each sample was extracted with a DNA extraction kit and using the field-ready Edwards buffer. B) Results using spores from *C. ribicola*. Spores were pooled from multiple blisters. Four extractions were performed and tested, two using a DNA extraction kit and two using direct PCR. C) Results obtained using a portable real-time PCR instrument. Two DNA extractions were prepared in the instrument and tested using lyophilized reagents (Extraction #2 and #3). DNA extracted using a kit was used as a positive control (Extraction #1).

### Shelf-life test of the field-ready lyophilized kits

As the lyophilized mastermix needs to be prepared and aliquoted in advance for use in the field, a crucial aspect of the usefulness of this approach is the long term stability of the kit without refrigeration. We prepared two batches of real-time PCR strips containing all the PCR reagents for the *S. musiva* assays, with and without trehalose and tested them at regular intervals. The strips were stored at room temperature in the dark and tested over the course of a year, running each assay in triplicate. We did not find a significant difference between the reactions with and without trehalose (F = 1.457, p>0.05) (Fig 2B). However, the reactions conducted without trehalose yielded no amplification after 263 days at room temperature. By comparison, amplifications were detected for all reactions with added trehalose until the end of the test, on day 389. The trehalose-added reactions had a small but significant effect age effect (F = 4.356, p<0.05), but no effect on the consistency of the replicates (F = 4.334, P>0.05); this could indicate a slight performance degradation over time. However, there were no false negatives in the reactions with trehalose at any time point.

## Real-time PCR field kit demonstration

To demonstrate the workability of our protocols in field applications, we combined both the field-ready lyophilized reagents and DNA extraction on a portable instrument without the use of additional lab equipment (see Standard Operating Procedure for point-of-use real-time PCR in the Supplementary Material). In the presented test, we used fixed volume, disposable, 20 µL pastettes, pre-aliquoted reagents in disposable containers and the lyophilised *Cronartium* reagents to perform real-time PCR in a portable Franklin real-time PCR unit (Biomeme), an instrument capable of performing 9 concurrent reactions and monitor each on 3 channels, including FAM and CY5, allowing the duplexing of two assays (Fig 1H). Spores of *C. ribicola* were prepared as described earlier. Two extractions were performed using the Franklin as a dryblock, and the real-time PCR reactions were conducted in duplicate with lyophilized reagents. DNA extracted using a column was used as a positive control. Both direct extractions resulted in a positive amplification, with C_t_ values for the direct extractions between 32 and 33.5 (Fig 3C, Table S8). Testing was also conducted on a two3 instrument (Fig 1G) (Biomeme) using the *S. musiva* and the Lymantria assays with similar results (results not shown). All the material and the instrument can be carried in a small backpack, demonstrating the potential of performing simple, cost-effective and user-friendly molecular testing in the field, at the point-of-use.

## Discussion

Forestry and environmental applications often require sampling in remote areas far away from laboratory facilities. The need for on-site DNA testing capability is growing in areas such as forest health monitoring, where detection of potentially invasive species requires immediate actions. In this proof-of-concept study, we demonstrated the transfer of existing real-time PCR assays for simple, easy-to-use, cost-effective, sample-to-data, point-of-use, real-time PCR identification. Although assays conducted in the laboratory with fresh reagents and sample grinding were more sensitive, the iSPEED assays did not yield false negatives. We do not envision that field-testing will replace lab-testing but instead provide a useful and efficient complementary on-site screening. Ultimately, once an undesired species is identified in the field, additional sampling and testing would be required.

Other portable assays have been developed for plant pathogens. LAMP assays are popular for field applications and allow rapid and sensitive detection of plant pathogens [47] and insects [48]. There are three advantages of the iSPEED protocols: 1) the ability to leverage the vast library of existing published and validated real-time PCR assays, 2) the extensive ecosystem supporting real-time PCR: multiple enzymes and chemistry, numerous types of assays and a growing range of portable instruments, and 3) cost-effectiveness. We demonstrated the broad applicability of the iSPEED protocols: they can be used on different types of material, ranging from infected deciduous tree leaves, evergreen perennial Rhododendron leaves, fungal rust spores and legs and antennae of adult moths. Further testing will be required for more recalcitrant material. For example, those containing wood with extractives will be more challenging and might require adjustments to the current protocols.

Although we developed our protocols with forest pests and pathogens applications in mind, they could also be applied to detect the presence of other invasive species, such as the American bullfrog (*Lithobates catesbeianus*) or the Red-eared slider (*Trachemys scripta elegans*) by quickly detecting their DNA from water samples. They could also be used to correctly identify fish sold at retail stores and restaurants, as it is frequently mislabeled (Willette et al., 2017). Finally, there are also potential medical applications, for example, thedetection of food borne pathogens like *Clostridium perfrigens*, for which a large number of real-time PCR assays have been developed (Albini et al., 2008; Chon et al., 2012; Fukushima et al., 2007; Gurjar et al., 2008; Shannon et al., 2007; Wise and Siragusa, 2005).

Future work should focus on improving and expanding the scope of available assays. Preliminary testing indicates real-time PCR mastermixes from manufacturers other than those of the Qiagen company used here are also suitable. The DNA extraction step could also be improved to increase sensitivity or expand the range of material suitable for testing. For example, the use of Chelex 100 resin or simple disruption tools such as micro-pestles and/or abrasives could be tested to improve yield. Also, more capable portable real-time PCR instruments are in development, allowing the development of more complex assays through multiplexing as well as providing wells for more samples to be tested per run.

## Authors contribution

**AC** Conceptualization, Investigation, Validation, Visualization, Writing – Original Draft Preparation

**KH** Investigation, Writing – Review & Editing

**KA** Resources

**MC** Resources, Writing – Review & Editing

**DS** Resources, Writing – Review & Editing

**GB** Resources, Writing – Review & Editing

**PT** Resources, Writing – Review & Editing

**NF** Resources, Writing – Review & Editing

**RH** Conceptualization, Writing – Original Draft Preparation

## Acknowledgments

The authors would like to thank Xianya “Sabrina” Qu for testing the protocol and Karen McLachlan Hamilton for providing the pheromone traps.

This work was funded by Genome Canada, Genome British Columbia, Genome Quebec, the Canadian Forest Service and the Canadian Food Inspection Agency, through a Genomics Applications Partnership Program (GAPP 6102; Genome Canada) grant. Also this work is funded by Genome Canada, Genome British Columbia, Genome Quebec, the Canadian Forest Service, the Canadian Food Inspection Agency and FP Innovations through a Large Scale Applied Research Project (LSARP 10106; Genome Canada) grant.

